# *Drosophila suzukii* wing spot size is robust to developmental temperature

**DOI:** 10.1101/800417

**Authors:** Ceferino Varón-González, Antoine Fraimout, Vincent Debat

**Author notes:** Contact information: Ceferino Varón González, Institut de Systématique, Evolution, Biodiversité (ISYEB), Muséum National d’Histoire Naturelle, CNRS, Sorbonne Université, EPHE, Université des Antilles, 57 rue Cuvier, CP 50, 75005 Paris, France.

## Abstract

Phenotypic plasticity is an important mechanism allowing adaptation to new environments and as such it has been suggested to facilitate biological invasions. Under this assumption, invasive populations are predicted to exhibit stronger plastic responses than native populations. *Drosophila suzukii* is an invasive species whose males harbor a spot on the wing tip. In this study, by manipulating developmental temperature, we compare the phenotypic plasticity of wing spot size of two invasive populations with that of a native population. We then compare the results with data obtained from wild-caught flies from different natural populations. While both wing size and spot size are plastic to temperature, no difference in plasticity was detected between native and invasive populations, rejecting the hypothesis of a role of the wing-spot plasticity in the invasion success. In contrast we observed a remarkable stability in the spot-to-wing ratio across temperatures, as well as among geographic populations. This stability suggests either that the spot relative size is under stabilizing selection, or that its variation might be constrained by a tight developmental correlation between spot size and wing size. Our data show that this correlation was lost at high temperature, leading to an increased variation in the relative spot size, particularly marked in the two invasive populations. This suggests (i) that *D. suzukii*’s development is impaired by hot temperatures, in agreement with the cold-adapted status of this species; (ii) that the spot size can be decoupled from wing size, rejecting the hypothesis of an absolute constraint and suggesting that the wing color pattern might be under stabilizing (sexual) selection; (iii) that such sexual selection might be relaxed in the invasive populations. Finally, a subtle but consistent directional asymmetry in spot size was detected in favor of the right side in all populations and temperatures, possibly indicative of a lateralized sexual behavior.

## Introduction

Phenotypic plasticity often plays an important role in the adaptation to new environments (Lande, 2015; West-Eberhard, 1989). In particular, it has been repeatedly suggested to facilitate invasions (Chown et al., 2007; Gibert et al., 2016; Richards et al., 2006), as genetic variation tends to be limited during the first stages of colonization (Geng et al., 2007; Geng et al., 2016). However, this possibility has received little empirical support so far (Arnold et al., 2019; Fox Rebecca J. et al., 2019). Invasions provide a particularly favorable context for the study of the evolutionary role of plasticity as we can compare derived populations repeatedly confronted to new environments with populations from the native range, arguably closer from the ancestral state. Differences in reaction norms are expected across populations (Pigliucci, 2005): native populations being predicted to be overall less plastic than invasive populations (Davidson et al., 2011; Lande, 2015; Lee and Gelembiuk, 2008; Richards et al., 2006; Yeh and Price, 2004).

Biological invasions are processes in which an increasing number of individuals from a population colonize one environment where no representative of that population was present. The population effective size fluctuates in the new habitat and the population phenotype may change during invasions (Travis et al., 2009), as natural selection (in a different environment) would gain importance relative to drift. The limited population size during colonization may increase the importance of genetic drift and decrease selection on sexual traits (Candolin and Heuschele, 2008; Lahti et al., 2009). Such relaxed sexual selection might in turn be associated with an increased variability of such traits (Marshall et al., 2008), and in particular an increased plasticity (Price, 2006).

*Drosophila suzukii* has recently colonized all continents in a wide and fast dissemination all over the world (Fraimout et al., 2017). This species is a particularly suitable model to investigate the role of plasticity in the success of an invasion. Remarkably, *D. suzukii* males present a dark spot on the wing, as in several other Drosophila species (Kopp and True 2002). A precise understanding of the effect of this spot on mating success is lacking (Fuyama, 1979; Roy and Gleason, 2019), but the particular courtship behavior (the male confronts the female showing her his dorsal wing area) consistently associated with the presence of a spot in males across species, suggests an important role in sexual selection (Kopp and True, 2002).

The effect of sexual selection on phenotypic plasticity is controversial (Greenfield and Rodriguez, 2004; Møller and Alatalo, 1999; Rowe and Houle, 1996). Although sexually selected traits are often considered to be plastic (Price, 2006), an increased robustness might evolve through stabilizing or directional selection (see Fierst (2013) for a theoretical treatment, and Nieberding et al. (2018) for a case study in *Bicyclus anynana*). Stabilizing selection would limit variation around the values preferred by females (Mead and Arnold, 2004), while directional selection would limit variation on one side of the distribution due to female preferences and on the other side due to developmental or physiological constraints (Wiernasz, 1989). We reasoned that *D. suzukii*’s spot might be more plastic, or at least less canalized and thus more variable in the invasive populations compared to native populations, owing to a hypothetical reduced choosiness in females – and thus reduced sexual selection – possibly advantageous in small, peripheral populations (Bleu et al., 2012).

Here we quantify the phenotypic variation of the wing spot size in several natural populations of *D. suzukii* and assess its plasticity to temperature using samples from three geographic populations reared in the lab. Our experimental samples allow us to propose a causal link between temperature and phenotypic variation, while our natural populations would confirm the existence of such an association in nature. Temperature is one of the main environmental drivers of life history and morphological evolution of drosophilids in particular and ectotherms in general (Atkinson, 1994; Crill et al., 1996; David et al., 1997; Gibert et al., 1996; Gibert et al., 2007) and the thermal plastic response in *D. suzukii* is receiving much attention (Clemente et al., 2018; Fraimout et al., 2018; Shearer et al., 2016). We test (1) whether natural populations present different spot sizes, and whether this differentiation correlates with local temperature at the time of capture, as would be expected for a plastic effect; (2) whether invasive and native populations display different plasticities to temperature; and (3) whether spot size is less canalized (more variable) in invasive populations. A higher plasticity of the spot in invasive populations is expected if plasticity plays a role in the invasion success. Combined with a reduced canalization, it might also indicate that sexual selection is relaxed in the invasive populations.

## Materials and Methods

### Samples

Natural populations were sampled in 2014 and 2016 in 13 localities worldwide (see table 1) with baited traps and net sweeping, and conserved in ethanol in the dark (Fraimout et al., 2017). The number of individuals per locality ranged from 13 to 50 (Table 1). Sample sizes correspond to samples and pictures originally collected to run morphometric analyses.

**Table 1.**
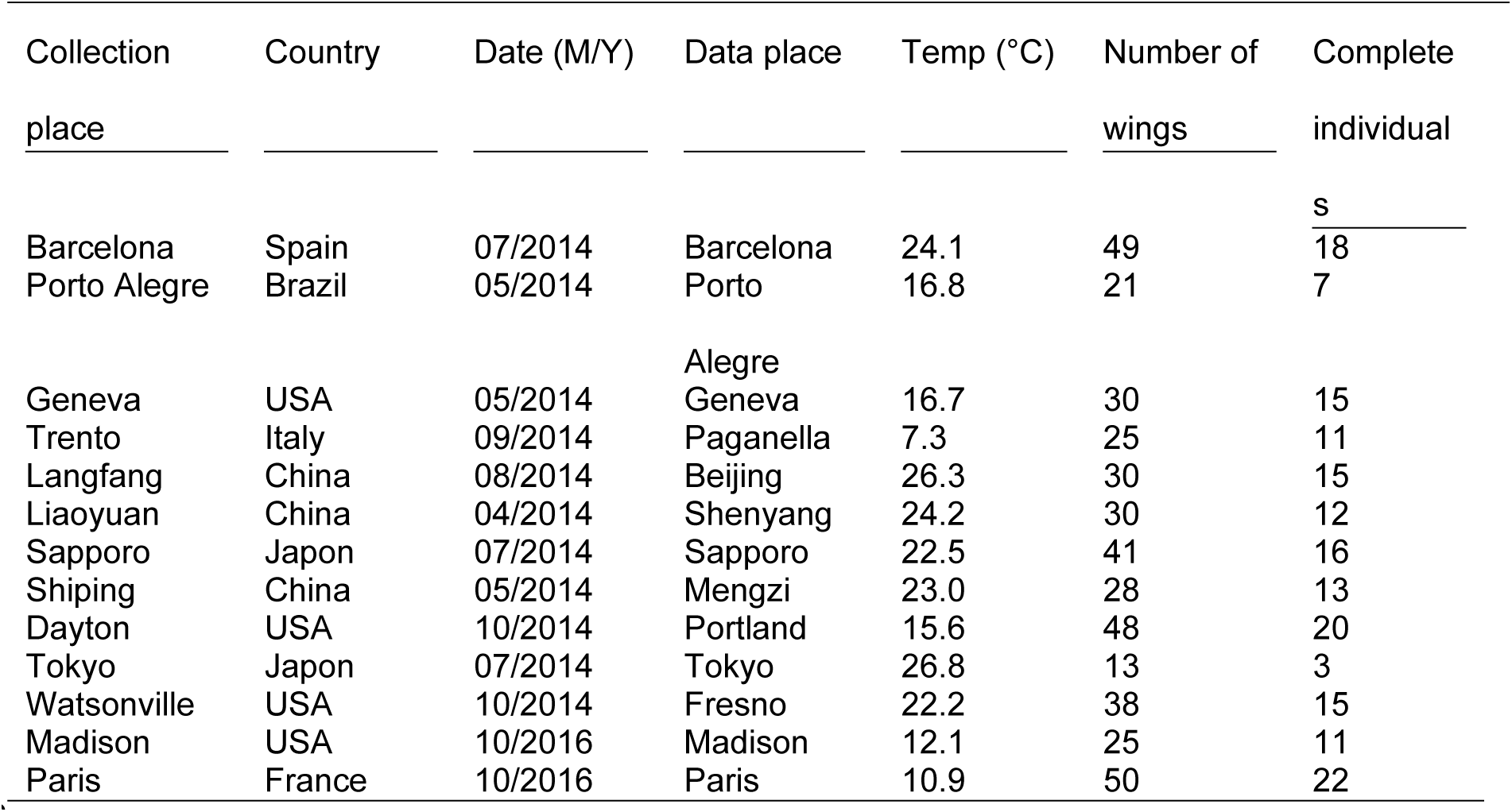
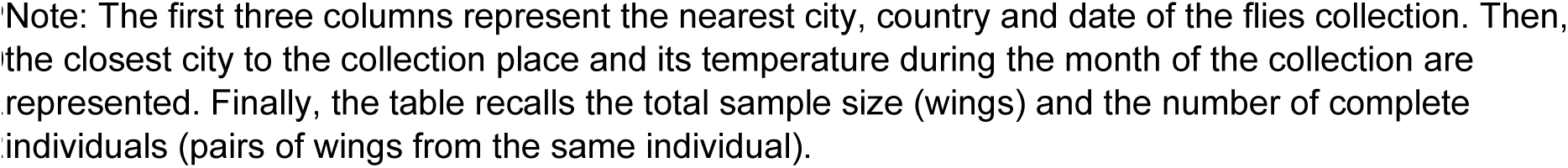
Natural populations sample sizes

| Collection<br>place | Country | Date (M/Y) | Data place | Temp (°C) | Number of<br>wings | Complete<br>individual<br>s |
| --- | --- | --- | --- | --- | --- | --- |
| Barcelona | Spain | 07/2014 | Barcelona | 24.1 | 49 | 18 |
| Porto Alegre | Brazil | 05/2014 | Porto Alegre | 16.8 | 21 | 7 |
| Geneva | USA | 05/2014 | Geneva | 16.7 | 30 | 15 |
| Trento | Italy | 09/2014 | Paganella | 7.3 | 25 | 11 |
| Langfang | China | 08/2014 | Beijing | 26.3 | 30 | 15 |
| Liaoyuan | China | 04/2014 | Shenyang | 24.2 | 30 | 12 |
| Sapporo | Japon | 07/2014 | Sapporo | 22.5 | 41 | 16 |
| Shiping | China | 05/2014 | Mengzi | 23.0 | 28 | 13 |
| Dayton | USA | 10/2014 | Portland | 15.6 | 48 | 20 |
| Tokyo | Japon | 07/2014 | Tokyo | 26.8 | 13 | 3 |
| Watsonville | USA | 10/2014 | Fresno | 22.2 | 38 | 15 |
| Madison | USA | 10/2016 | Madison | 12.1 | 25 | 11 |
| Paris | France | 10/2016 | Paris | 10.9 | 50 | 22 |

549Note: The first three columns represent the nearest city, country and date of the flies collection. Then, 550the closest city to the collection place and its temperature during the month of the collection are 551represented. Finally, the table recalls the total sample size (wings) and the number of complete 552individuals (pairs of wings from the same individual).

The laboratory lines used here were adopted from an earlier study on *D. suzukii’*s wing plasticity to developmental temperature (see Fraimout *et al.* 2018 for details on rearing and maintenance). Briefly, three natural populations of *D. suzukii were* sampled in 2014 with banana bait traps and net swiping: one in the native area (Sapporo, Japan), and two in the invasion range, in Paris (France) and Dayton (Oregon, USA). Ten isofemale lines per population were produced from single matings of random pairs of individuals performed in separate vials (David et al., 1997). For each line from each population, we transferred F1 full-siblings individuals in multiple replicated vials and repeated the process for five generations in order to amplify stock populations prior to the temperature treatment. Stock populations were kept at 22°C throughout the experiment.

At the onset of the temperature treatment, we sampled random individuals from each line and transferred them in two sets of new vials. After 24h – and once oviposition was ensured – adult flies were removed and one set of vials was placed in an incubator set at 16°C, while the other set was placed in an incubator at 28°C. All vials were randomly placed within incubators and kept at each experimental temperature for the full development of the flies (i.e. egg to adult). The remaining temperature of our treatment (i.e. 22°C) corresponded to the one used for stock populations.

Between eight and 17 individuals per line were obtained (Table 2). Due to limitation on sample sizes, we did not take into account the lines in the analyses. We mounted wings on slides in a mixture of ethanol and glycerin and then we sealed the coverslips with nail polish and small weights to keep the wings as flat as possible. Images were taken with a Leica DFC 420 digital camera mounted on a Leica Z6 APO microscope controlling for the scale.

**Table 2.** Lab populations sample sizes

| Geographic origin | Temperature (°C) | Number of wings | Complete<br>individuals | Lines |
| --- | --- | --- | --- | --- |
| Paris | 16 | 15 | 7 | 8 |
|  | 22 | 27 | 12 | 15 |
|  | 28 | 28 | 13 | 15 |
| Sapporo | 16 | 25 | 12 | 13 |
|  | 22 | 26 | 12 | 14 |
|  | 28 | 31 | 15 | 16 |
| Dayton | 16 | 22 | 10 | 12 |
|  | 22 | 26 | 11 | 15 |
|  | 28 | 31 | 14 | 17 |

557Note: The first two columns show the geographic origin of each population (first) and the temperature 558set during their development *in-vitro* (second). Then, the total number of wings used to analyze overall 559variation and the number of complete individuals used to analyze asymmetric variation.

### Phenotypic data

Two phenotypic traits were directly inferred from the pictures: wing size and spot size (Figure 1). Because the potential visual effect of the wing spot during courtship may be associated to the proportion of the wing occupied by the spot and not its absolute size, we also estimated the relative spot size as the ratio between spot size and wing size for each individual. Using the ratio between spot and wing size for each individual can lead to erroneous conclusions due to allometry between traits (e. g. Packard and Boardman, 1988) but the estimation of residual values from a linear regression between wing and spot size gave virtually identical results (see online data).

**Figure 1:**
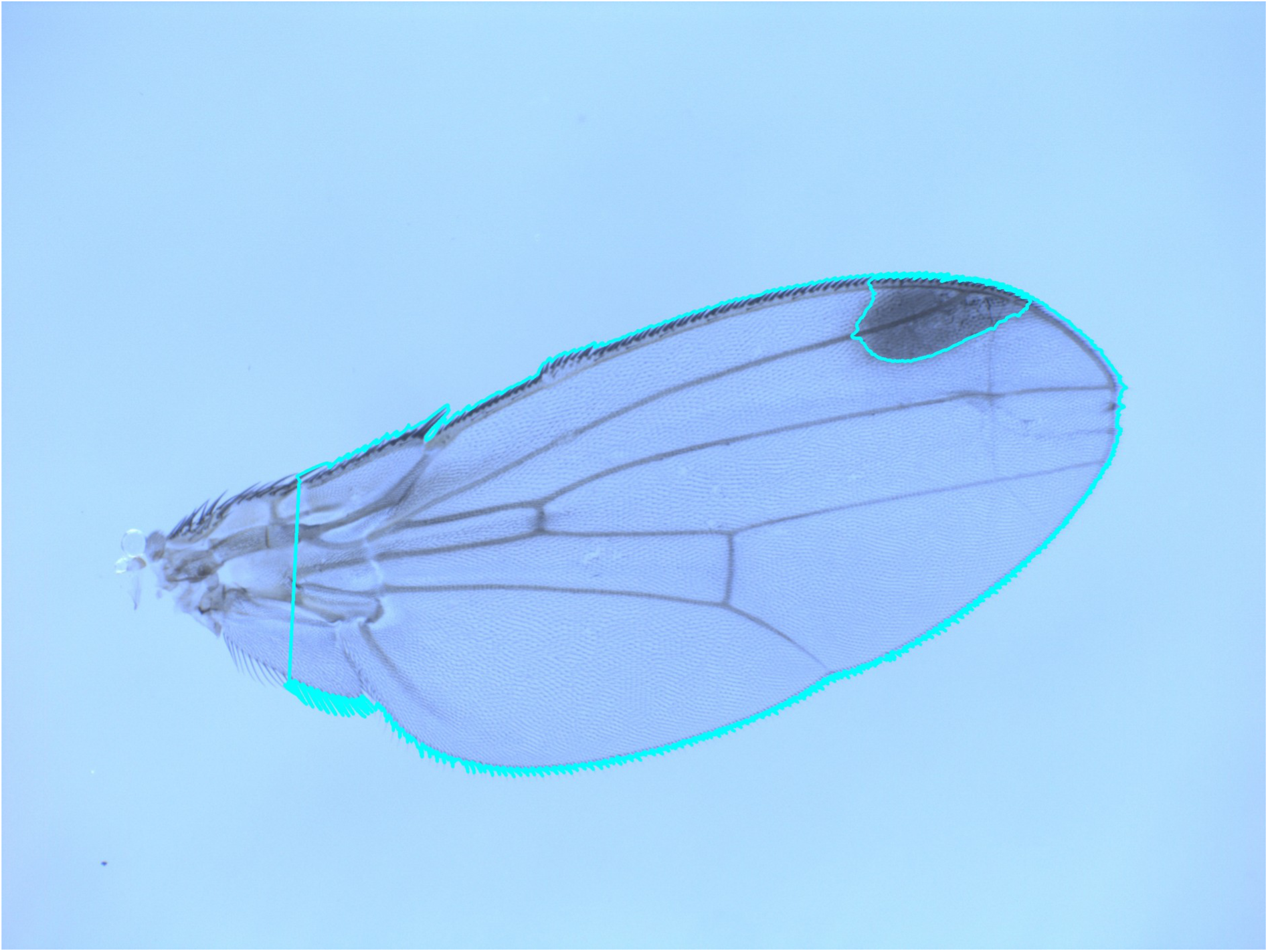
*Drosophila suzukii* wing from a population of Sapporo (Japan, native population) raised at 28°. In blue there is the wing and spot areas identified by our Python algorithm.

In order to remove the proximal part of the wing, which is usually deteriorated during the removal of the wing from the thorax and may bias wing-size estimates, two landmarks were placed in ImageJ 1.51j8 (Rasband, 2012), that correspond to landmarks 1 and 3 in Fraimout et al. (2018) and a line between them was traced to discard all the proximal part of the wing. Then, a semi-automatized script in Python 3.6.1 was written to extract the wing and spot contours. Their respective areas were estimated by the Green’s theorem in the function contourArea of the OpenCV library (Bradski and Kaehler, 2000) as the number of pixels within the wing and spot contours.

Our script was based on the manual adjustment of the picture color contrast. Because the color difference between the wing and the background is sharp, the wing outline was easily and robustly extracted. To the contrary, because the limits of the spot are fuzzy, a subtle difference in the image contrast can induce a relatively large difference in the spot outline. The manual adjustment of the picture contrast might thus be an important source of measurement error, particularly so for the spot size. We thus repeated the estimation of the wing and spot sizes in all the pictures and measurement error was quantified using an ANOVA for each population, assessing individual variation and measurement residuals.

### Statistical analyses

Natural populations - We applied a linear model to assess the relationship between absolute spot size and wing size, differences among natural populations on the spot size and whether there was natural variation in the spot-wing relationship (i. e. spot size ~ wing size + population + wing size x population). To have an estimate of natural variation for the other two phenotypic traits, we also assessed with one-way ANOVAs the differences among natural populations for wing size and the ratio. To explore the potential effect of temperature on those wild samples, we collected the average temperature during the month of collection in these places from the National Oceanic and Atmospheric Administration database (www.ncdc.noaa.gov/IPS/mcdw/mcdw.html) (Table 1) and ran a linear regression for each trait with temperature as a continuous variable.

Experimental samples - We inferred the effect of temperature and geographic factors on each phenotypic trait with two-way ANOVAs (phenotype ~ temperature + population + temperature x population). The interaction temperature x population was used to assess the difference in plasticity among populations. We also ran a regression to test the effect of wing size on the absolute spot size, so that we can experimentally develop a null hypothesis about the relationship between wing and spot sizes that can be compared later with the results from natural populations. Because temperature, wing size and spot size are correlated (see results), we did not include two of these factors as explanatory variables in the same statistical model to avoid collinearity problems. This is the simplest approach that allows us to explore the unique contributions of temperature and wing variation to spot variation (Graham, 2003). The effect of temperature on the relationship between wing size and spot size was assessed by estimating only the effect of the interaction wing size x temperature on spot size in one ANCOVA. Differences in the slope of the relationship spot size-wing size among temperature levels with three pairwise t-test.

Differences in variance might reflect differences in the developmental channeling of environmental variation (Debat and David, 2001; Debat et al., 2009). In general, stressful environmental conditions have been suggested to alter developmental stability, leading to an increase in variance among individuals and asymmetry (Graham et al. 1998).

Among individual variation was measured as the coefficient of variation (CV) to account for a potential scaling effect on wing and spot size variation, and compared among temperatures within each geographic population. The significance of the differences was tested using a modified signed-likelihood ratio (MSLR) test for equality of coefficients of variation (Krishnamoorthy and Lee, 2014).For all the analyses described we averaged the phenotypic measurements obtained over the two wings within individual to avoid pseudo-replication.

To explore the hypothesis that temperature affects asymmetry, we measured the difference between right and left values for each of the three phenotypic traits (Table 1). That way, positive values in asymmetry would reflect larger phenotypes in the right wing and negative values larger phenotypes in the left wing. Then, a two-way mixed model ANOVA was applied to each trait (Palmer and Strobeck, 1986; Palmer 1994). “Individual” was entered into the model as a random effect, and tested for the variation among individuals; “side” was treated as a fixed effect, and tested for a systematic deviation from perfect symmetry (directional asymmetry, DA); the interaction “side x individual” tested for the significance of non-DA relative to measurement error (Fluctuating asymmetry, FA). The replicated measurements were included in the error term. Once the presence or absence of significant DA and FA were assessed in each particular group, we assessed the effect of temperature and geography on these two traits with an ANOVA (effect on DA). Differences in the slope of the relationship temperature-ratio asymmetry among populations with three pairwise t-test. Finally, Levene’s tests were applied on the asymmetry measures to assess FA.

As effect sizes, Cohen’s *d* for differences among geographic populations and *r*^*2*^ for temperature effects were reported due to their simplicity (Rosenthal et al., 1994). All analyses were run in R version 3.5.2 (R Core Team 2013). All data, scripts and figures can be found in Dryad and Github.

## Results

### Natural populations

In natural populations, we found differences among populations for the spot size (ANOVA: F_12, 223_ = 2.480, *p* = 0.005) as well as for the wing (ANOVA: F_12, 236_ = 41.330, *p* < 0.001) and the ratio (ANOVA: F_12, 236_ = 3.092, *p* < 0.001). The latter showed a remarkable stability in comparison to the wing and the absolute spot size (*r*^*2*^_*RATIO*_ = 0.14 *r*^*2*^_*SPOT*_ = 0.49, *r*^*2*^_*WING_POP*_ = 0.68).

Our linear model (spot size ~ wing size + population + wing size x population) also showed an association between spot size and wing size in pooled samples (*r*^*2*^ = 0.66, F_1, 223_ = 525.217, *p* < 0.001) (Figure 2). Whether the relationship between spot size and wing size depends on the population is not conclusive (*r*^*2*^ = 0.70, ANOVA: F_12, 223_ = 1.748, *p* = 0.058). We detected a significant effect of temperature on the three phenotypic traits (Figure 2), but much weaker for the ratio (wing size: *r*^*2*^ = 0.39, F_1, 247_ = 158.1, *p* < 0.001; spot size: *r*^*2*^ = 0.21,F_1, 247_ = 68.88, *p* < 0.001; ratio: *r*^*2*^ = 0.03, F_1, 247_ = 8.183, *p* = 0.005).

**Figure 2:**
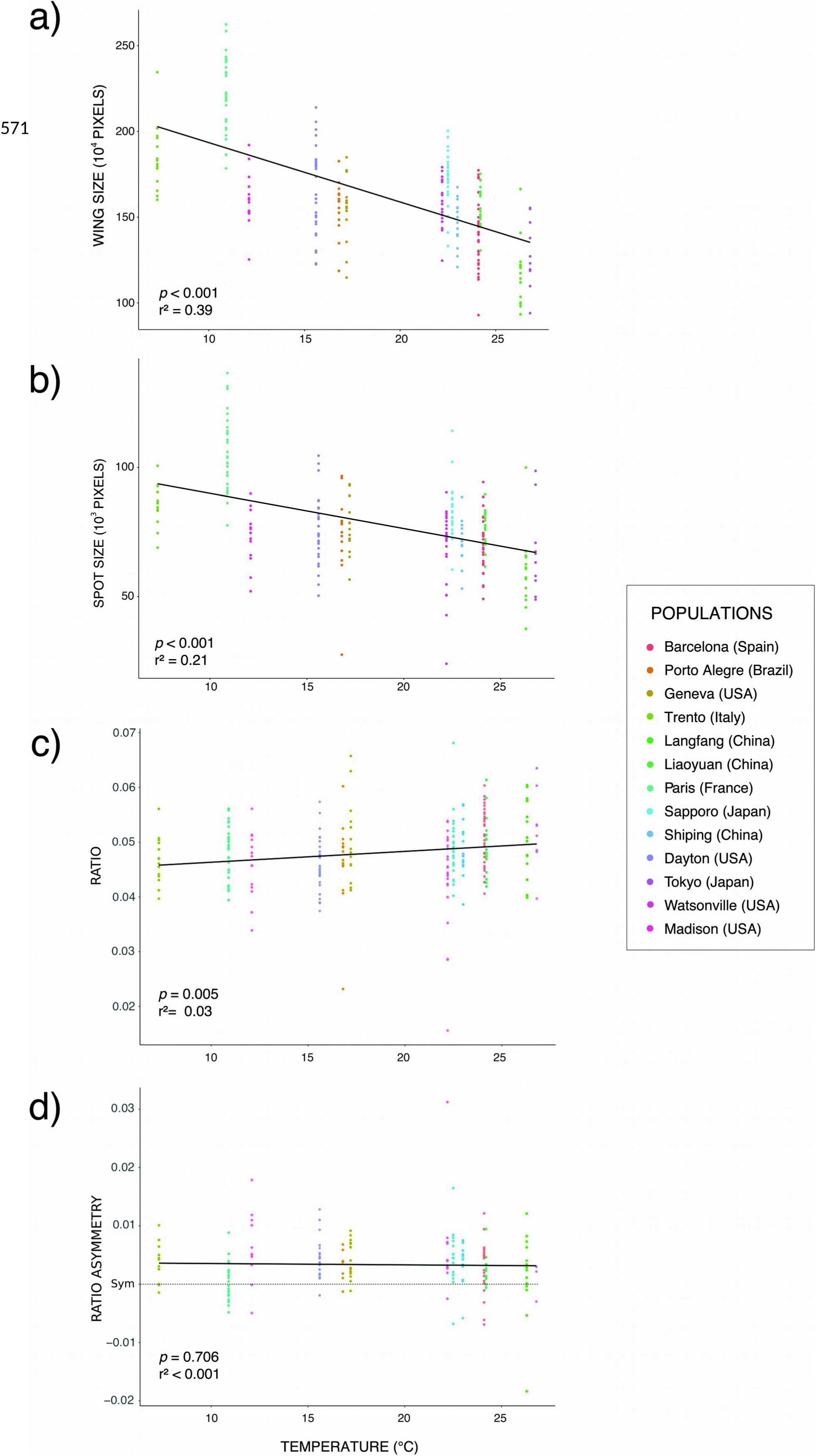
Relationship between wing and spot variation in natural populations and temperature. Temperature was measured in °C and phenotypic sizes correspond to the number of pixels within wing and spot areas in the pictures. The ratio and ratio asymmetry are based on a proportion between the former two. For relationships between the variables stated in the diagram axes, *p* and r^2^ were reported. Association between a) wing size and temperature, b) spot size and temperature, c) between the ratio and temperature and d) ratio asymmetry and temperature.

### Experimental samples

Both wing size and spot size strongly decreased with temperature (*r*^*2*^_*WING_TEMP*_ = 0.93, F_2, 116_ = 886.278, *p* < 0.001; *r*^*2*^_*SPOT_TEMP*_ = 0.67, F_2, 121_ = 4.986, p = 0.008) (Figure 3) and both traits are associated (*r*^*2*^ = 0.73, *t* = 18.017, *p* < 0.001) (Figure 3). The ratio exhibited much less variation with temperature (*r*^*2*^ = 0.08, F_2, 116_ = 4.907, *p* = 0.009), the values at 22°C being slightly larger and driving the significance (no difference between 16 and 28°C was detected: F_1, 75_ = 1.243, *p* = 0.268).

**Figure 3:**
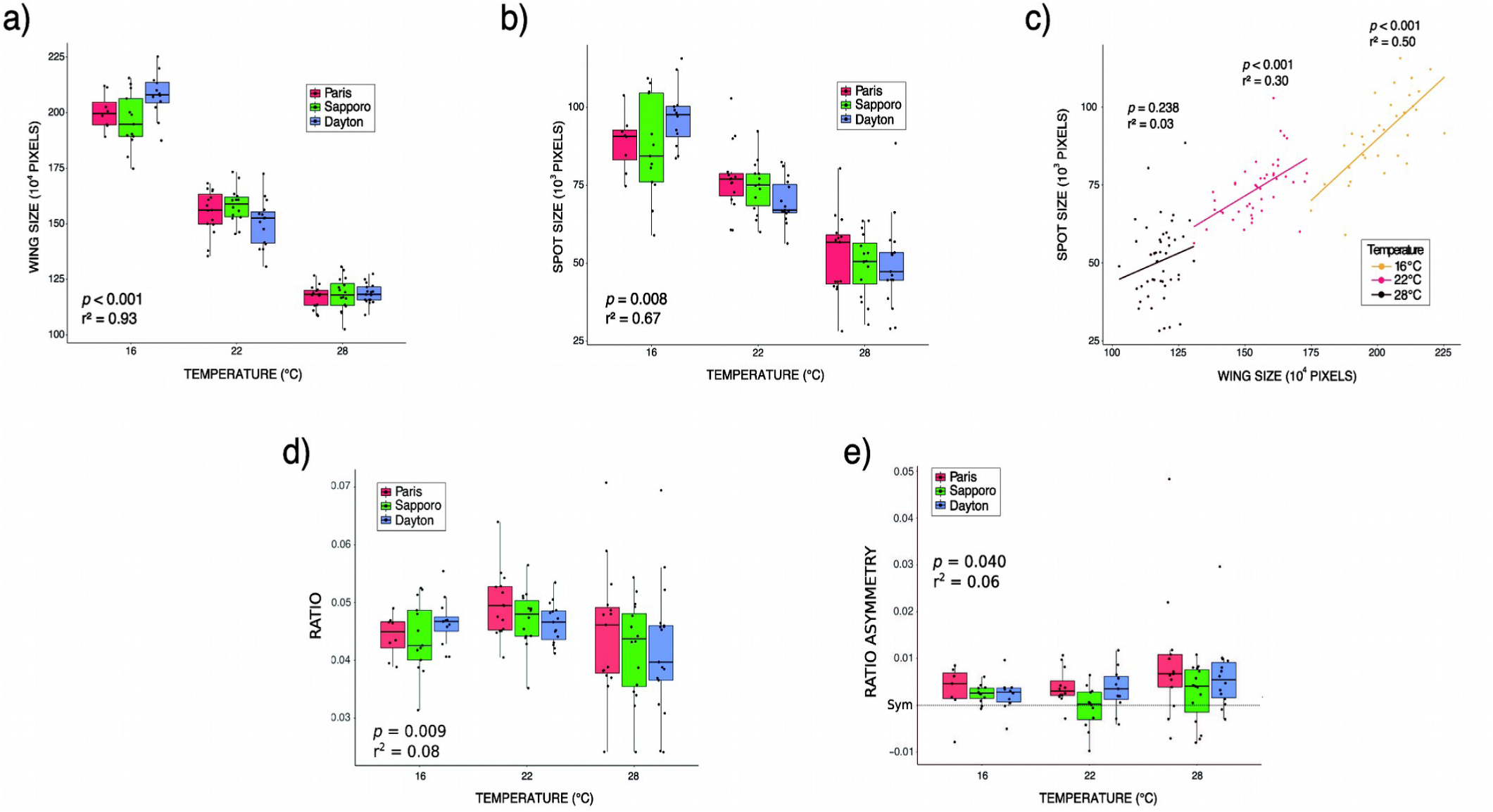
Effect of temperature on the different phenotypic traits in experimental samples. Temperature was measured in °C and phenotypic sizes correspond to the number of pixels within wing and spot areas in the pictures. The ratio and ratio asymmetry are based on a proportion between the former two. For relationships between the variables stated in the diagram axes, *p* and r^2^ were reported. a) linear regression showing the influence of developmental temperature on wing variation, b) linear regression showing the influence of developmental temperature on spot size variation, c) effect of temperature on the relationship spot size – wing size, d) differences in the ratio for each population among temperatures, e) differences in ratio asymmetry for each population among temperatures.

Overall, the three traits exhibited very similar reaction norms across populations (Figure 3a and 3c). A slight difference was nevertheless detected for wing size plasticity (interaction temperature x population significant: F_4, 116_ = 4.366, *p* = 0.003), the Sapporo population being slightly less responsive to temperature (*r*^*2*^_*SAPPORO*_ = 0.92, *r*^*2*^_*PARIS*_ = 0.94, *r*^*2*^_*DAYTON*_ = 0.95). No difference in plasticity among populations was found for the either the wing spot (F_4, 116_ = 1.928, *p* = 0.110) or the ratio (F_, 116_ = 0.508, *p* = 0.730).

Interestingly, the relationship between absolute spot size and wing size was dependent on temperature (*r*^*2*^ = 0.74, F_3, 121_ = 117.4, *p* < 0.001): the lower the temperature, the higher the slope of the regression and the tighter the association (*r*^*2*^_16°_ = 0.50, *r*^*2*^_22°_ = 0.30, *r*^*2*^_28°_ = 0.03) (Figure 3b). Although no significant differences were found among slopes (16°C vs 22°C: t_1, 73_ = −1.472, *p* = 0.145, 28°C vs 22°C: t_1, 88_ = −0.417, *p* = 0.678, 28°C vs 16°C: t_1, 77_ = −1.206, *p* = 0.232), this may be due to the sample size limitations within temperature.

Within temperature variation tended to be higher at 28°C for both the spot size and the ratio, although Sapporo did not show significant differences among temperatures for any of the phenotypic traits (MSLRT_SPOT_ = 4.25, *p* = 0.119, MSLRT_WING_ = 0.83, *p* = 0.661, MSLRT_RATIO_ = 4.60, *p* = 0.100). For absolute spot size, variation was higher at 28°C in comparison to 22°C in all three geographic populations: Sapporo (Table 3), Dayton (Table 3, MSLRT = 12.01, *p* < 0.001) and Paris (Table 3, MSLRT = 3.42, *p* = 0.064). At 16°C, the other extreme temperature, Sapporo population also showed increased variation in the spot size (Table 3) but Paris (Table 3, MSLRT = 1.01, *p* = 0.315) and Dayton did not (Table 3, MSLRT = 0.06, *p* = 0.807). For the ratio, although all three geographic populations showed increased variation at 28° compared to 22° (Table 3), Sapporo showed reduced variability (Table 3) F_2, 40_ = 1.02, *p* = 0.369). This might be explained by the fact that Sapporo showed very small wing variation among temperatures (Table 3; MSLRT = 0.83, *p* = 0.661).

**Table 3.** Coefficient of variation for each trait per population and temperature

|  |  | Sapporo | Dayton | Paris |
| --- | --- | --- | --- | --- |
| **Wing** | 16° | 0.06 | 0.05 | 0.04 |
|  | 22° | 0.05 | 0.07 | 0.06 |
|  | 28° | 0.06 | 0.04* | 0.04 |
| **Spot** | 16° | 0.19 | 0.10* | 0.10* |
|  | 22° | 0.11 | 0.10* | 0.14** |
|  | 28° | 0.20 | 0.29 | 0.25 |
| **Ratio** | 16° | 0.14* | 0.09* | 0.08* |
|  | 22° | 0.11* | 0.08* | 0.12* |
|  | 28° | 0.20 | 0.27 | 0.25 |

563Note: Asterisks represents a significant difference for a phenotype (\*: $\alpha = 0.05$ , \*\*: $\alpha = 0.1$ ), within a
564population, between the temperature presenting the largest coefficient of variation and the other two
565temperatures.

Measurement error was much lower than the FA detected for all the populations and traits, ranging from twice lower for wing size in Dayton at 28°C to near 200 times lower for spot size in Paris at 28°. No difference in FA was detected for either the wing or the absolute spot sizes for laboratory samples (F_WING; 2, 103_ = 2.449, *p* = 0.091; F_SPOT; 2, 103_ = 1.166, *p* = 0.316) and for natural populations no difference in FA was found for any trait (F_WING; 12, 165_ = 0.942, *p* = 0.507; F_SPOT; 12, 165_ = 0.597, *p* = 0.843; F_RATIO; 12, 165_ = 0.712, *p* = 0.417). For the ratio, the populations at 28° showed more FA than the populations at 16° (SD_RATIO 28°_: 0.010, SD_RATIO 16°_: 0.003, F_1, 69_ = 6.070, *p* = 0.016) but the differences between 22° and 16° (SD_RATIO 22°_: 0.005, F_1, 62_ = 2.069, *p* = 0.155) and 28° and 22° (F_1, 75_ = 3.543, *p* = 0.064) were not significant.

No DA was detected for wing size. In contrast, significant DA was detected for both absolute and ratio, always in favor of the right side. For Paris and Dayton, DA was detected at 22 and 28°C while Sapporo only showed significant DA at 16°C. DA subtly increased with temperature for the ratio (r^2^ = 0.06, F_2, 97_ = 3.323, *p* = 0.040) and this effect was different depending on the population (r^2^_TEMP x POP_ = 0.15, F_8, 97_ = 2.095, *p* = 0.043). In particular, native population showed a lower response to temperature than invasive populations (Figure 3d, *d*_*SAPPORO-PARIS*_ = 0.60, *d*_*SAPPORO-DAYTON*_ = 0.49, *d*_*DAYTON-PARIS*_ = 0.21), although pairwise tests did not show significant differences in the slope of the relationship temperature-ratio asymmetry between Sapporo with either Paris (t_1,65_ = 0.765, *p* = 0.447) or Dayton (t_1,68_ = 1.268, *p* = 0.209).

Most of the natural populations showed similar asymmetry patterns: while no DA was detected for wing size, both the absolute spot size and the ratio exhibited right biased DAs (Figure 2). The exceptions are the Dayton population, which also showed DA for the wing size, Langfang, which did not show DA for either the wing or the spot size, and Paris, which only showed wing size DA. DA was not associated with temperature in any trait (Wing: F_1, 165_ = 0.090, *p* = 0.765; absolute spot size (F_1, 165_= 0.488, *p* = 0.486); ratio (F_1, 165_ = 0.143, *p* = 0.706); Figure 2).

## Discussion

Our results revealed the existence of a relatively large natural variation in wing and spot sizes and temperature may explain a substantial part of it. In contrast, the spot-to-wing ratio was quite stable in nature and very robust to developmental temperature in the lab. This stability might either reflect a tight structural connection between the development of the spot and that of the wing, imposing a strong correlation (i.e. constraint hypothesis), or a history of stabilizing selection on the relative spot size (i.e. adaptive hypothesis). All three populations exhibited an increase in relative spot size variation at 28°C, indicative of a lesser developmental robustness at high temperature, in agreement with the status of cold-adapted species of *D. suzukii* (Enriquez et al., 2018; Jakobs et al., 2015; Shearer et al., 2016; Stephens et al., 2015). This increased variation originating from a de-correlation of spot size and wing size at 28°C suggests that the stability of the relative spot size is not an absolute structural constraint, and therefore points at the adaptive hypothesis. *D. suzukii* males wing color pattern might thus be under stabilizing (sexual?) selection. Native and invasive populations showed very similar reaction norms, suggesting that spot plasticity is not affected by – and played no role in the success of – the invasion history. Relative spot size variability was higher in invasive populations compared to the native population, possibly pointing at a relaxed sexual selection during the invasion. Finally, both natural and laboratory samples exhibited a slight but significant DA of spot size (both absolute and relative), consistently in favor of the right side.

### Relative spot size robustness: selection or constraint?

Although our experimental populations show a slight but significant decrease in the ratio at both high and low temperatures, the overall pattern reflects a global stability to developmental temperature, contrasting with both wing size and the absolute spot size that are both strongly plastic to temperature.

Such robustness might suggest either that spot size is fully determined by wing size, owing to the tight connection between wing development and the spot formation – an example of developmental constraint – or alternatively, that stabilizing selection is acting on *D. suzukii* males color pattern, removing individuals with extreme relative spot sizes.

The male-limited presence of the wing spot in all species has been interpreted as a mark of sexual selection (Kopp and True 2002). The presence of a spot is indeed phylogenetically tightly associated with the occurrence of elaborate courtship behaviors, the males dancing in a stereotypic way in front of females and displaying their wings (Kopp and True, 2002; Revadi et al., 2015). Although the empirical evidence of an effect of the spot on mating success is scarce (Fuyama, 1979; Roy and Gleason, 2019), it is nevertheless conceivable that females choice might be influenced by the male wing phenotype. Sexual selection might have favored the stability of the relative spot size and reduced its plasticity, a common process (Fierst, 2013) although not universal (Price, 2006). This would suggest that the focal trait might not be the spot size itself, but rather its extent on the wing, i.e. the wing color pattern.

Alternatively, it is possible that spot size plasticity is simply a structural consequence of wing size plasticity to temperature, which has been suggested to be adaptive (Crill et al., 1996; David et al., 1997), and the fact that the boundaries of the expression of the genes involved in the spot formation vary according to the positional information of the veins (Arnoult et al., 2013; Gompel et al., 2005). Such developmental dependency of spot formation on wing development would thus impose a tight correlation between spot size and wing size, leading to a structurally stable spot relative size. This would thus be an example of an absolute constraint on the variation of the wing color pattern.

The analysis of the ratio variation within temperature provides an element in the discussion. It was indeed found that this variation increases at high temperature in the three populations (Figure 3c). This effect may indicate that hot temperatures might destabilize the processes involved in the spot formation, which is in agreement with the idea that *D. suzukii* is adapted to cold temperatures (Jakobs et al. 2015; Stephens et al. 2015; Shearer et al. 2016; Enriquez et al. 2018). The analysis of the covariation between wing size and spot size shows that they are tightly correlated. However, this correlation decreases at 28°C overall (Figure 3b), leading to the observed increase in variation of the ratio. This shows that the developmental link between wing and spot development can be disrupted (e.g. under hot temperature) suggesting that the stability of the spot relative size under colder (possibly optimal) temperatures might reflect more than a developmental constraint: i.e. might be maintained by stabilizing selection.

### No role for plasticity to temperature in the invasion

Because plasticity may give a fitness advantage in early stages of invasion, populations repeatedly confronted to changing environments as invasive populations might present larger plastic responses. The hypothesis of adaptive plasticity in invasive populations (as opposed to native ones) has frequently appeared in the literature (Davidson et al., 2011; Lande, 2015; Lee and Gelembiuk, 2008; Richards et al., 2006; Yeh and Price, 2004). However, our results only show very small differences in wing size plasticity between invasive and native populations.

These results are consistent with some of the previous studies on the invasion of *D. suzukii.* No difference in thermal plasticity of wing shape, activity rhythms, gene expression and ovipositor shape was detected (Fraimout et al., 2018; Plantamp et al., 2019; Varón-González et al., 2019). There is is still scarce evidence that thermal plasticity might have contributed to the success of the invasion in this species (but see e. g. Everman et al. 2018). It is possible that *D. suzukii* might have been preadapted in its native range to the thermal conditions encountered throughout the invasion (Suarez and Tsuitsui, 2008), reducing the importance of temperature as a selective agent, but leaving open the hypothesis that plasticity relative to other parameters might have played a role in the invasion success (Hamby et al., 2016).

### Directional asymmetry: affected by heat stress and indicative of a lateralized sexual behavior?

Both DA and FA have been suggested to increase under stress and have in turn been used as bioindicators of stressful environmental conditions (Graham et al., 1998; Klingenberg and Nijhout, 1999), but see Houle (1998) for a critical discussion). Our results, showing an increase of the spot DA at high temperature may be circumstantial but their congruence with independent suggestions about the adaptation of *D. suzukii* populations to cold temperature (Enriquez et al., 2018; Jakobs et al., 2015; Shearer et al., 2016; Stephens et al., 2015) might advise further exploration.

The systematic DA in favor of the right side for the spot size is surprising. The small magnitude of DA might be a consequence of a regular but non adaptive effect (Pélabon and Hansen, 2008). However, this consistent right-biased DA possibly points at a bias during courtship and mating that might be worth investigating, as suggested by QTL analyses linking behavior and wing pigmentation in *Drosophila elegans* and *Drosophila gunungcola* (Yeh and True, 2014). By altering a trait involved in a sexual behavior, stressful developmental conditions (temperature) might thus interfere with sexual selection.

## Acknowledgements

We would like to thank K. Tamura, M. Toda, P. Shearer, S. Fellous and T. Schlenke for their help during the fly sampling. We also thank M. Guillaume for her help with the fly stock maintenance and F. Peronnet for the rearing medium used. We thank Jennifer Gleason for her words on the reproduction of *D. suzukii* and the potential role of directional asymmetry on it. We also thank Thomas Flatt and two anonymous reviewers for the comments in a previous version of this article. CVG would like to thank Hacklab Almeria for their comments on the Python algorithm to estimate the wing and spot areas. CVG and AF were funded by the Agence Nationale de la Recherche (ANR) under the project SWING (ANR-16-CE02-0015).

## Data accessibility

All data, R scripts and reports of R scripts can be found in Dryad.

## Competing interests

The authors declare no competing or financial interests.

## Author contributions

Conceptualization: VD and CVG. Flies collection: AF and VD. Experimentation: AF. Statistical analysis: CVG. Discussion and writing: CVG and VD.

## Notes

#### Summary of Updates

Substantial changes in the statistical analyses and the results, which did not affect in a significant way the discussion and conclusion of the study

